# Screening of CHO-K1 endogenous promoters for expressing recombinant proteins in mammalian cell cultures

**DOI:** 10.1101/2021.10.13.464282

**Authors:** Ileana Tossolini, Agustina Gugliotta, Fernando López Díaz, Ricardo Kratje, Claudio Prieto

**Affiliations:** UNL, CONICET, FBCB (School of Biochemistry and Biological Sciences), CBL (Biotechnological Center of Litoral), Ciudad Universitaria, Ruta Nacional 168 Km 472.4, C.C. 242. (S3000ZAA), Santa Fe, Argentina; UNL, FBCB (School of Biochemistry and Biological Sciences), CBL (Biotechnological Center of Litoral), Ciudad Universitaria, Ruta Nacional 168 Km 472.4, C.C. 242. (S3000ZAA), Santa Fe, Argentina; Cellargen Biotech S.R.L., Antonia Godoy 6369, CP 3000, Santa Fe, Argentina; Regulatory Biology Laboratory, Salk Institute for Biological Studies, 10010 N. Torrey Pines Road, La Jolla, CA 92037

**Keywords:** CHO cells, Endogenous promoters, Mammalian cells, Transcriptional activity

## Abstract

For the production of recombinant protein therapeutics in mammalian cells, a high rate of gene expression is desired and hence strong viral-derived promoters are commonly used. However, they usually induce cellular stress and can be susceptible to epigenetic silencing. Endogenous promoters, which coordinates their activity with cellular and bioprocess dynamics while at the same time they maintain high expression levels, may help to avoid such drawbacks. In this work, endogenous promoters were identified based on high expression levels in RNA-seq data of CHO-K1 cells cultured in high density. The promoters of Actb, Ctsz, Hmox1, Hspa5, Vim and Rps18 genes were selected for generating new expression vectors for the production of recombinant proteins in mammalian cells. The *in silico*-derived promoter regions were experimentally verified and the majority showed transcriptional activity comparable or higher than CMV. Also, stable expression following a reduction of culture temperature was investigated. The characterized endogenous promoters (excluding Rps18) constitute a promising alternative to CMV promoter due to their high strength, long-term expression stability and integration into the regulatory network of the host cell. These promoters may also comprise an initial panel for designing cell engineering strategies and synthetic promoters, as well as for industrial cell line development.

## 1. Introduction

The production of recombinant proteins is one of the most important areas in the biopharmaceutical industry. Chinese hamster ovary (CHO) cells are the most widely used platform for the industrial production of therapeutic proteins. ^[1]–[3]^ Other mammalian cell lines thar are commonly used include BHK-21 cells and HEK293T cells. ^[2], [4]^ Cell line development is continuously improving towards increased productivities, faster production, and process development. But, the generation of high-producing cell lines is still a laborious and multi-step process. ^[5]^ Many technologies for optimization of heterologous protein production have been developed. However, the most pivotal control mechanism, the transgene expression promoter, has often been ignored.

The choice of a suitable promoter is a crucial stage to achieve very high and stable transgene expression in mammalian cell cultures. ^[6]^ Today, viral-derived promoters, such as human cytomegalovirus (hCMV) or simian virus 40 (SV40), are frequently used for constitutive expression of recombinant proteins in mammalian cells. On one hand these promoters allow high expression rates, but on the other hand they can lead to constitutive overexpression of the transgene causing a permanent stress on the cell ^[6]^. In addition, viral promoters show cell-cycle dependency ^[7], [8]^. The CMV, for example, is active primarily in the S phase and presents very low activity in G0/G1 ^[8]^. In a previous work, we found a G0/G1 synchronization mainly in late exponential and stationary growth phases in CHO-K1 cells cultured in high density. ^[9]^ This is particularly important because the G1-phase is considered the appropriate stage for increasing the production of recombinant proteins. ^[10]^ Moreover, several studies showed that CMV promoter could be susceptible to histone modifications ^[11], [12]^ or DNA methylation, ^[13]–[18]^ linking these epigenetic modifications to production instability in recombinant CHO cells.

A solution to such drawbacks would be the use of endogenous promoters which function in a coordinated manner with cellular and bioprocess characteristics. Some cellular promoters have been investigated for this purpose in the last few years, for example, by using libraries with several genomic fragments, or flanking sequences of highly expressed housekeeping genes, by analyzing transcriptome microarray or RNA-seq data. ^[19]–[24]^ However, few analyses have been done considering the expression stability during long-term culture of those promoters.

In this study, endogenous promoters were identified from RNA-Seq data of CHO-K1 cells cultured in high density. ^[9]^ Promoters of genes with high levels of expression in both, late exponential and stationary growth phases, were selected as potential candidates for industrial application. The promoter sequences of Actb, Ctsz, Hmox1, Hspa5, Vim and Rps18 genes were used to generate new expression vectors. These vectors proved to be useful for high-level protein expression not only in CHO, but also in BHK and HEK cells. The majority of these endogenous promoters constitute an alternative to CMV promoter, and they are meant to achieve high expression level, long-term expression stability and improve production process efficiency.

## 2. Materials and Methods

### 2.1. Screening of novel-promoters activity in mammalian cells

#### 2.1.1. Selection of candidate endogenous promoters and Generation of expression vectors

Candidate promoters were selected based on RNA-seq data of CHO-K1 suspension-adapted cells cultured in 1 L bioreactors under a temperature gradient from 37 °C to 31 °C. ^[9]^ Genes with high levels of expression in both exponential and stationary phases were chosen based on their FPKM values. DNA regions upstream of highly expressed genes were visualized on IGV tool ^[25]^ using CHO-K1 genome reference and mapped reads, to detect potential transcription start sites (TSS), promoter location and enhancer regions. Also, a comparison with promoters of homologous genes (*M. musculus, R. norvegicus* and *H. sapiens*) was performed. These sequences were retrieved from Eukaryotic Promoter Database (EPD). ^[26]^ A subset of 50 genes that occurred in all datasets was then selected to form a ‘high transcriptional activity group’ (Table S1, Supporting Information). From that group, six potential promoter candidates were randomly selected for subsequent cloning, characterization and testing for promoter functionality *in vitro*.

The endogenous promoters were generated by PCR from genomic DNA of suspension adapted-CHO-K1 cells. Primers were designed using Primer3 ^[27]^ and primer-BLAST ^[28]^ in order to amplify 900-1000 bp upstream the TSS (Table S2, Supporting Information). To achieve directional cloning, SacI and KpnI restriction sites were introduced at 5’ ends of primers forward and reverse, respectively. PCR products were digested with SacI and KpnI (Promega) and ligated in the pZsGreen1-1 plasmid, using a standard T4 DNA ligase protocol (Invitrogen). The pZsGreen1-1 vector lacks eukaryotic promoter and enhancer sequences, and carries the coding sequence of the green fluorescent protein ZsGreen1, derived from *Zoanthus sp*. Also, it has a cassette containing a neomycin resistance gene, for stable integration in mammalian cells. The sequences of all plasmid constructs were confirmed by dideoxy DNA sequencing (Macrogen).

#### 2.1.2. Cell cultures

CHO-K1 cells were cultured in adherent conditions in chemically defined medium DMEM-F12 (Gibco), supplemented with 5% (v/v) of fetal bovine serum (FBS) (PAA Laboratories), 200 mM L-glutamine (Sigma-Aldrich), 2.441 g/L NaHCO_3_ (Gibco) and 0.05 mg/mL gentamicin sulfate (Gibco). HEK293T cells were maintained in 13.37 g/L DMEM medium (Gibco), supplemented with 10% (v/v) of FBS (PAA Laboratories), 1.5 g/L NaHCO_3_ (Gibco), 0.05 g/L gentamicin sulfate (Gibco) and 0.11 g/L sodium pyruvate. BHK-21 cells were cultured in MEM medium (Gibco), supplemented with 10% (v/v) FBS (PAA Laboratories), 2.2 g/L NaHCO_3_ (Gibco) and 0.05 g/L gentamicin sulfate (Gibco). Cells lines were cultured in a humidified atmosphere at 37 °C and 5% CO_2_.

#### 2.1.3. Transient expression

CHO-K1, BHK-21 and HEK293T cells were transfected with 1 µg of novel vectors containing the endogenous promoters. The pCMV-ZsGreen1-1 was used as control to compare the transcriptional activity of endogenous promoters. Co-transfection of each plasmid with pCIneo-CMV-IFNα vector ^[29]^ was performed to evaluate transfection efficiency, determined by quantification of interferon alpha (IFNα) in culture supernatants. At the same time, novel vectors were co-transfected with pBKS plasmid used as a carrier to equalize DNA load across all transfections. Lipofectamine^*TM*^ 2000 Transfection Reagent (Invitrogen) was used, according to the manufacturer’s instructions.

After 48 h, transfection efficiency was analyzed by assessing fluorescence intensity of ZsGreen1 in transfected cells by fluorescence microscopy (Eclipse Ti-S, Nikon Instruments Inc) and flow cytometry (Guava EasyCyte, Merck Millipore). The percentage of ZsGreen1-expressing cells (%ZsGreen) and ZsGreen1 mean fluorescence intensity (MFI) of each sample were analyzed using Guava ExpressPlus software (Merck Millipore) and MFI was expressed as fluorescence arbitrary units (FAU). Additionally, the culture supernatant of pCIneo-CMV-IFNα co-transfected cells was collected and analyzed by ELISA sandwich using the protocol described by Ceaglio *et al*. (2008). ^[29]^

#### 2.1.4. Generation of stable pools

CHO-K1 and BHK-21 cells were transfected with plasmid constructions (as described above) and then cultured under neomycin (NEO) (Gibco) selection pressure. Resistant cells were grown in culture medium without selection pressure to investigate the stability of the transgene. The stably transfected cells were screened for ZsGreen1 expression during several generations. Analyses were performed by flow cytometry.

#### 2.1.5. Effect of temperature shift on levels of stable ZsGreen1 expression in CHO-K1 cells

CHO-K1 stable cell lines were cultured in two replicates in 24-well plates with a cell density of 2.0×10^5^ cells/mL at generation 25. The plates were maintained for 10 days at 37 °C (control cultures) and at 32 °C, in a humidified atmosphere and 5% CO_2_. ZsGreen1 expression was analyzed by flow cytometry on days 3, 5, 7 and 10. The MFI of ZsGreen1 positive cells for each promoter was plotted.

#### 2.1.6. Statistical analysis

Transient transfections were performed in duplicates. Differences were assessed using one-way ANOVA and two different post hoc tests: Tukey’s test (for multiple comparison) and Dunnett’s test (for comparing with a control sample).

In expression stability studies, the results obtained during the whole experiment were analyzed by one-way ANOVA followed by Tukey’s and Dunnett’s post hoc tests. The transcriptional activity of endogenous promoters at 32 °C and 37 °C was analyzed at four different time points (3, 5, 7 and 10 days) in duplicate. These results were analyzed by a Student’s t-test.

Parameters were considered significantly different when p < 0.05. GraphPad Prism version 6.00 (GraphPad Software Inc.) was used to perform data analysis and graphing.

### 2.2. In silico analysis of endogenous promoter sequences

Potential transcription factor binding sites (TFBSs) were identified using Alibaba2, ^[30]^ applying a minimum matrix conservation value of 80%; Match, ^[31]^ using a vertebrate matrix and a cut-off value set to minimize false positive matches; and GPMiner, ^[32]^ setting default values against mouse database. A search for putative CpG islands was carried out using the EMBOSS cpgplot software. ^[33]^ In addition, promoter conservation for orthologous genes from *M. musculus, R. norvegicus* and *H. sapiens*, was analyzed. Promoter sequences were compared by pairwise global sequence alignment using the EMBOSS needle software. ^[33]^

## 3. Results

### 3.1. Selection of promoter regions based on RNA-seq of CHO-K1 suspension cell cultures

Selected endogenous promoters were identified starting from RNA-Seq data previously generated. ^[9]^ These data were obtained from three replicates of CHO-K1 suspension-adapted cells that were cultured in 1 L bioreactors. Transcriptional levels of all genes were ranked from high to low, based on their FPKM values. A subset of genes from each phase was selected in order to identify those that were highly active at late exponential and stationary growth phases. Finally, six potential candidate promoters (not previously described in other studies) were selected for subsequent cloning, characterization and testing of promoter functionality *in vitro*: Actb, Ctsz, Hmox1, Hspa5, Rps18 and Vim.

### 3.2. Screening of transcriptional activity of endogenous promoters in mammalian cells

Promoter regions of selected genes were PCR-amplified from genomic DNA of suspension-adapted CHO-K1 cells and then cloned into pZsGreen1-1 vector.

Despite many authors have demonstrated that EF1-alpha is a useful promoter for stable and transient transgene expression, some of them evidenced no significant differences when comparing with CMV. ^[34],[35]^ Particularly, Mufarrege *et al*. (2014) ^[36]^ have shown that stable CHO cell lines producing recombinant human Factor VIII, under the control of either CMV or EF1-alpha, presented no significant differences in their productivities. For this reason, we decided to employ the pCMV-ZsGreen1-1 plasmid to compare the transcriptional activity of endogenous promoters with CMV. This viral promoter presents strong transcriptional activity in mammalian cells culture *in vitro* that allows high expression of transgenes. Such vector contains the CMV sequence, the immediate-early enhancer/promoter region, a chimeric intron and the coding sequence of ZsGreen1.

#### 3.2.1. Transfection efficiency and transient transgene expression

Firstly, a transient transfection assay was performed to evaluate transcriptional activity of isolated endogenous promoters, not only in CHO-K1 but also in HEK293T and BHK-21 cells. Transient gene expression is particularly useful in early research or preclinical studies when several potential therapeutic candidates are needed for evaluation or when a molecule is required in the short term. ^[37], [38]^

After 48 h, transfection efficiency was analyzed by quantifying IFNα concentration using an ELISA sandwich (data not shown). No significant differences between IFN concentrations among transfections was observed for each tested cell line (ANOVA test, P ≤ 0.05), thus transcriptional activity of endogenous promoters was evaluated by assessing fluorescence intensity of ZsGreen1 in transfected cells by fluorescence microscopy (Figure 1.A) and flow cytometry (Figure 1.B-G and Figure S1, Supporting Information). When comparing co-transfections of each plasmid containing the novel promoter, either with pBKS or pCIneo-CMV-IFNα, we could observe that ZsGreen1 expression was practically the same (Figure 1.B and C for CHO-K1 cells; Figure 1.D and E for HEK293T cells; Figure 1.F and G for BHK-21 cells). No expression of ZsGreen1 was determined using the wild type cell line (negative control) which was set as marker 1 (M1) in flow cytometry analysis. The expressing population was divided as cells with moderate fluorescence intensity (marker 2, M2, MFI between 10^1^ and 10^3^) and cells with high fluorescence intensity (marker 3, M3, MFI higher than 10^3^). Analyzing M3 is particularly interesting because promoters with high transcriptional activity are attractive candidates for driving transgene expression in mammalian cells.

**Figure 1.**
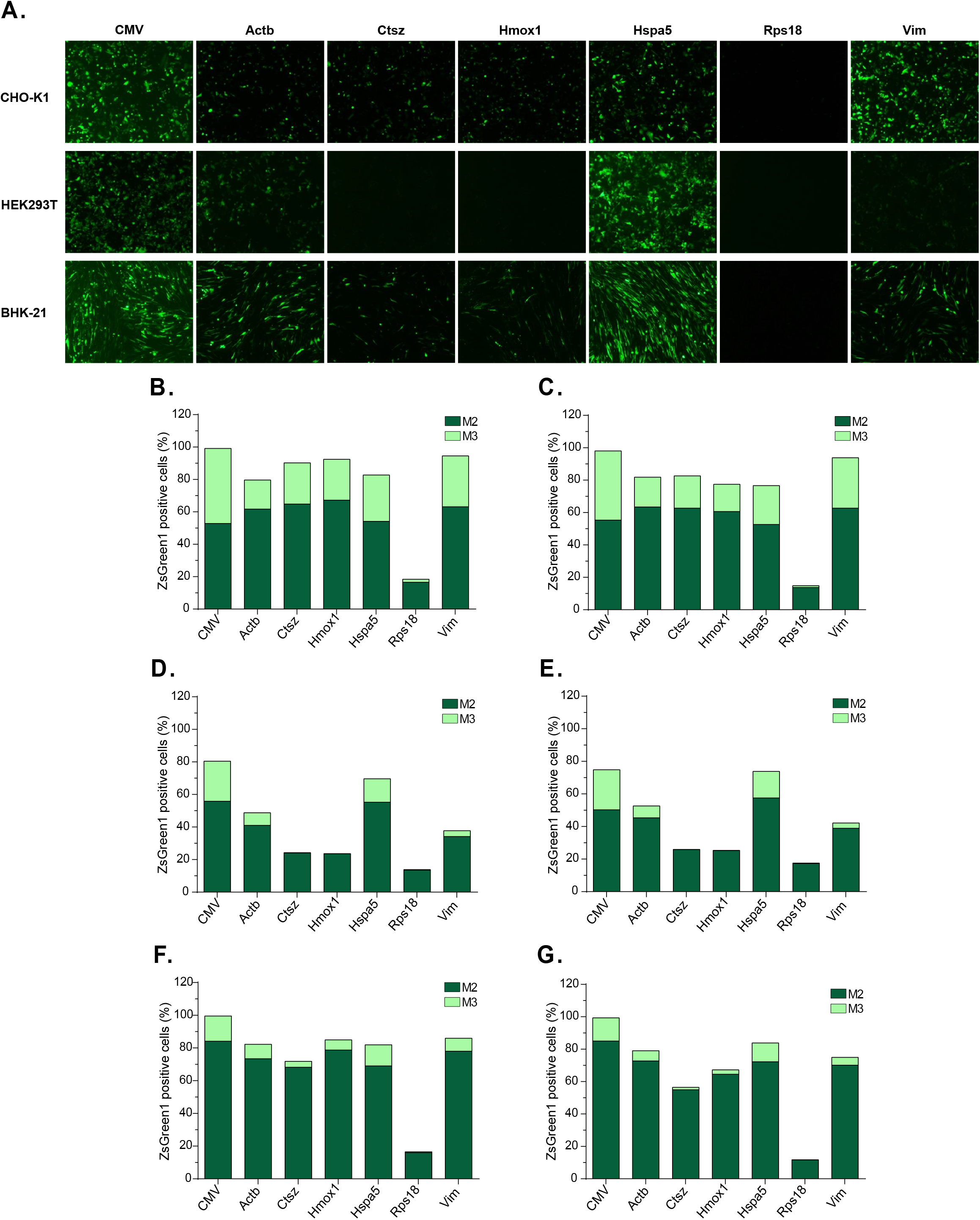
Effect of different promoters on transient transgene expression. pZsGreen1-1 vectors containing CMV, Actb, Ctsz, Hmox1, Hspa5, Rps18 and Vim promoters were transfected into CHO, HEK293T and BHK-21 cells. **(A)** ZsGreen1 fluorescence profile of transfected cells (100x, 200 ms exposure time). **(B, D, F)** %ZsGreen1 for CHO-K1, HEK293T and BHK-21 cells, respectively, co-transfected with new vectors carrying endogenous promoters and pBKS plasmid. **(C, E, G)** %ZsGreen1 for CHO-K1, HEK293T and BHK-21 cells, respectively, co-transfected with new vectors carrying endogenous promoters and pCIneo-CMV-IFNα plasmid. **M2:** positive cells with moderate fluorescence intensity, from 10^1^ to 10^3^ FAU (Fluorescence arbitrary units). **M3:** positive cells with high fluorescence intensity, FAU values higher than 10^3^.

Hamster-derived cell lines (BHK-21 and CHO-K1) exhibited enhanced transient transgene expression, and, as expected, CHO-K1 showed the highest ZsGreen1 expression levels. It is important to mention that Rps18 showed the lowest transcriptional activity in all cell lines (ANOVA test, P<0.05, Figure 1). For this reason, it was excluded from the subsequent analysis. This endogenous promoter exhibited high expression levels during suspension culture in bioreactor, ^[9]^ which might be attributed to the bioprocess dynamic and the assistance of additional regulatory elements for gene expression.

CHO-K1 cells transfected with all endogenous promoter-containing vectors displayed ∼80-90% of ZsGreen1 positive cells. Vim activity was comparable to CMV promoter since nearly 100% of the cells expressed ZsGreen1 (ANOVA test, P<0.05, Figure 1.B-C). In general, HEK293T cells exhibited the lowest transcriptional activity (significant differences were found when comparing transcriptional activity of each promoter in human and hamster-derived cells, ANOVA test P<0.05). This can be explained by the fact that promoters were isolated from a hamster genome. Particularly, CMV and Hspa5 showed around 70% of ZsGreen1 expressing cells (Figure 1.D-E), and the other promoters displayed inferior transcriptional activity (∼25-50%).

In BHK-21 cells (Figure 1.F-G), CMV displayed the highest percentage of Zsgreen1 positive cells (99%), while endogenous promoters Actb, Hmox1, Hspa5 and Vim showed ∼70-85% of cells expressing the transgene. However, no significant differences between the transcriptional activity of endogenous promoters and CMV were found. Ctzs exhibited ∼60% ZsGreen1 positive cells which was significantly lower compared to CMV (ANOVA test, P<0.05).

MFI analysis indicates that ZsGreen1 expression in CHO-K1 and BHK-21 cells was high (in M3) and practically constant (Figure S1.B, F, Supporting Information), showing values around 3.0-4.0×10^3^ FAU (fluorescence arbitrary units). In HEK293T cells, differences among MFI values were especially marked. In particular, CMV, Actb, Ctsz and Hspa5 promoters reached MFI values higher than 3.8×10^3^ FAU (Figure S1.D, Supporting Information).

#### 3.2.2. Stable levels of long-term transgene expression driven by endogenous promoters

Long-term stable expression of the transgene is crucial for the industrial production of recombinant proteins. As transient expression cannot completely reveal the function of different promoters, it is important that vectors can be stably expressed in cells, therefore we analyzed the stability of ZsGreen1 under the control of endogenous promoters in hamster-derived cell lines.

For cell line generation, CHO-K1 and BHK-21 cells were transfected with endogenous promoter-containing vectors or pCMV-ZsGreen1-1 plasmid, and selective pressure was performed using increasing concentrations of NEO. This process allowed the gradual selection of cells that expressed increasing ZsGreen1 intensity. BHK-21 cells resisted to higher concentrations of the antibiotic compared to CHO-K1 cells (5.0 vs. 4.0 mg/mL NEO, respectively). Cell lines selected with the maximum concentrations of NEO exhibited the highest expression of ZsGreen1 (Figure S2.A-B, Supporting Information). CHO-K1 and BHK-21 cell lines resistant to the maximum concentration of NEO were grown in culture medium without antibiotic, in order to analyze ZsGreen1 expression stability. After 120 h, the expression of the reporter protein remained practically constant for all promoters in both cell lines (Figure S2.C-D, Supporting Information), meaning that NEO is not necessary to maintain high expression levels.

The stably transfected cell lines were screened for ZsGreen1 expression during several generations (Figure 2 and Figure S3, Supporting Information). Figure 2.A shows the stability of ZsGreen1 expression in CHO-K1 cells during 45 generations. The %ZsGreen1 was notably higher for cell lines containing endogenous promoters compared to CMV (ANOVA test, P<0.05), except for Rps18 promoter that exhibited the lowest expression (ANOVA test, P<0.05). From now onwards this promoter (Rps18) will be excluded from the analysis. Cells carrying Hspa5 promoter showed the highest %ZsGreen1, which was nearly 100% during all generations. Actb, Vim and Hmox1 promoters exhibited similar transcriptional activity to Hspa5 (no significant differences were observed after ANOVA test and Dunnett’s post hoc test, p<0.05). In general, transcriptional activity of endogenous promoters remained relatively constant and stable during the 45 generations, while CMV activity varied periodically over time, from 40 to 80 %ZsGreen1. Even though CMV promoter delivers high gene expression, some works reported protein production instability. ^[17], [18], [39]–[41]^ Gene silencing mechanism of this promoter is not clear, although DNA methylation, histone hypoacetylation, activation of signaling molecules and altered levels of transcriptional factors have been suggested. ^[11]–[13], [15]–[17], [42]–[44]^

**Figure 2.**
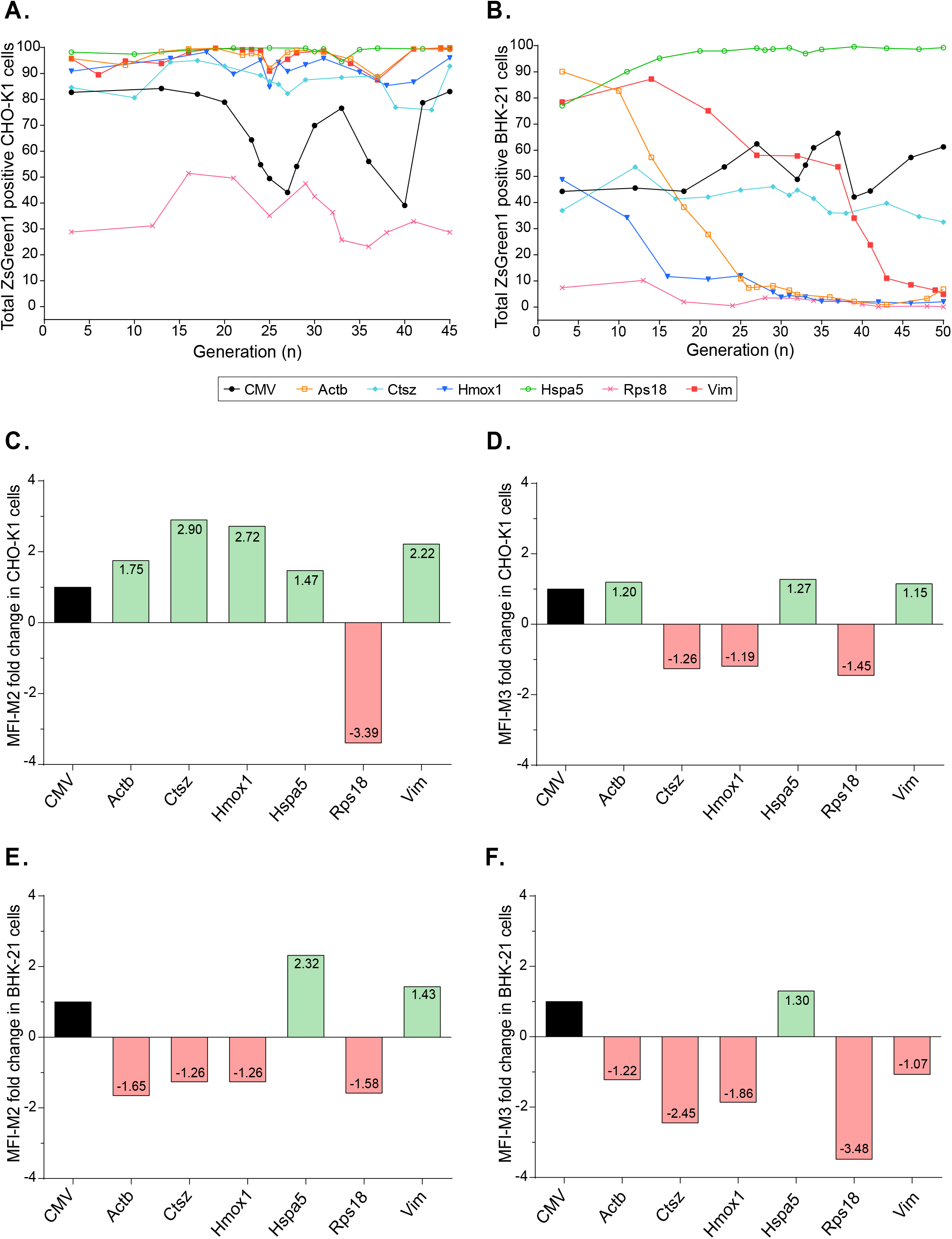
ZsGreen1 expression stability in CHO-K1 and BHK-21 cells lines grown until 45 and 50 generations, respectively, in the absence of neomycin selection pressure. Total %ZsGreen1 positive CHO-K1 and BHK-21 cells (**A** and **B**, respectively). Moderate ZsGreen1 MFI (M2) fold change in CHO-K1 and BHK1-21 cells (**C** and **E**, respectively). High ZsGreen1 MFI (M3) fold change in CHO-K1 and BHK-21 cells (**D** and **F**, respectively). **MFI:** Mean Fluorescence Intensity. **M2:** positive cells with moderate fluorescence intensity, from 10^1^ to 10^3^ FAU (Fluorescence arbitrary units). **M3:** positive cells with high fluorescence intensity, FAU values higher than 10^3^.

For CHO cell lines, the percentage of each population expressing moderate (M2) and high (M3) levels of ZsGreen1 during 45 generations, are shown in Figure S3.A-B (Supporting Information). The lowest % of cells expressing ZsGreen1 at high fluorescence intensity (M3) was observed for CMV promoter, whereas the highest (∼70%) was seen for Hspa5 promoter (ANOVA test, P<0.05). Actb and Vim promoters also exhibited a relatively high proportion of cells expressing ZsGreen1 with high intensity (∼50-70% in M3). However, no significant differences were found between them (ANOVA test, P<0.05). Furthermore, MFI values among generations were represented for each cell line, distinguishing between cells expressing ZsGreen1 in M2 and M3 (Figure S4, Supporting Information). These results demonstrated that all endogenous promoters presented higher ZsGreen1 intensity in M2 compared to CMV (Figure S4.A, Supporting Information).

In Figure 2.C-F, MFI is expressed as fold change, taking CMV as control (reference value as 1.0). In the case of CHO-K1 cell populations in M2 (Figure 2.C), Ctsz and Hmox1 showed the strongest promoter activity (2.90-fold and 2.72-fold, respectively), followed by Vim (2.22-fold), Actb (1.75-fold) and Hspa5 (1.47-fold). When analyzing CHO-K1 cells in M3 (Figure S4.B, Supporting Information), it was observed that Hspa5, Actb and Vim promoters drove ZsGreen1 expression with a clearly higher fluorescence intensity compared to CMV promoter; exhibiting fold-change values of 1.27, 1.20 and 1.15, respectively (Figure 2.D).

For BHK-21 cells, ZsGreen1 expression stability during 50 generations is shown in Figure 2.B. In general, less stable and non-uniform transcriptional activity was observed for BHK-21 cells compare to CHO-K1 cell line. Hspa5-ZsGreen1 cell line exhibited the highest percentage of positive cells, approximately 100% after 15 generations (ANOVA test, P<0.05). Also, Hspa5 allowed ZsGreen1 expression in a more stable manner than the rest of endogenous promoters. CMV activity exhibited a slight variation over time, and the proportion of cells expressing ZsGreen1 was near 50% (Figure 2.B). ZsGreen1 expression under the control of Actb and Hmox1 promoters notably decreased after 10 generations and in the case of Vim promoter, after 15 generations. Ctsz transcriptional activity was rather constant, but the percentage of positive cells was lower than the observed for CMV-ZsGreen1 cell line (t-test, p=0.0006).

The percentage of cells in M3 was extremely low for all BHK cell lines (Figure S3.D, Supporting Information). The only exception was Hspa5-promoter containing up to 70% of positive cells in M3. The results shown in Figure S4.C (Supporting Information) highlight Hspa5 as the promoter with the greatest activity in BHK-21 cells, since it exhibited the highest values of MFI, in M2. In terms of MFI fold-change, taking CMV as control with a reference value of 1.0 (Figure 2.E-F), only Hspa5 and Vim showed increased MFI for cells expressing ZsGreen1 in M2 (2.32 and 1.43-fold, respectively). In the case of cells in M3 maker, Hspa5 exhibited a fold change of 1.30.

#### 3.2.3. Effect of temperature shift on stable transgene expression

A temperature reduction step is commonly used during production cell culture in the biopharmaceutical industry. ^[45]–[51]^ The initial rapid cell growth phase is performed at 37 °C followed by a growth arrest phase when the culture is shifted to a lower temperature (typical 30 - 32 °C). This change in temperature is used for simultaneously inducing growth arrest and extending culture viability thus increasing recombinant protein productivity and yield.

ZsGreen1 stable expression, driven by endogenous promoters and CMV, was examined in CHO-K1 cells at 32 °C and 37 °C during 10 days (Figure 3). For temperature shift, 32 °C was selected because the endogenous promoters were identified based on RNA-seq data of CHO-K1 cells cultured at 33 - 31 °C during stationary phase. ^[9]^ Moreover, 32°C represents the average temperature applied by other studies mentioned before. Figure 3 shows that almost all promoters enhanced their activity at 32 °C. Student’s t-test revealed significant differences between ZsGreen1 positive cells at 37 and 32 °C for the following promoters: Actb, Ctsz and Rps18 (p=0.04, p=0.014, and p=0.01, respectively). In the case of Vim and Hmox1 promoters, significant differences were found when comparing % of cells in M3 (p=0.015 and p=0.014, respectively). No significant differences were observed for Hspa5. However, it was expected since this promoter evidenced more than 99% positive cells.

**Figure 3.**
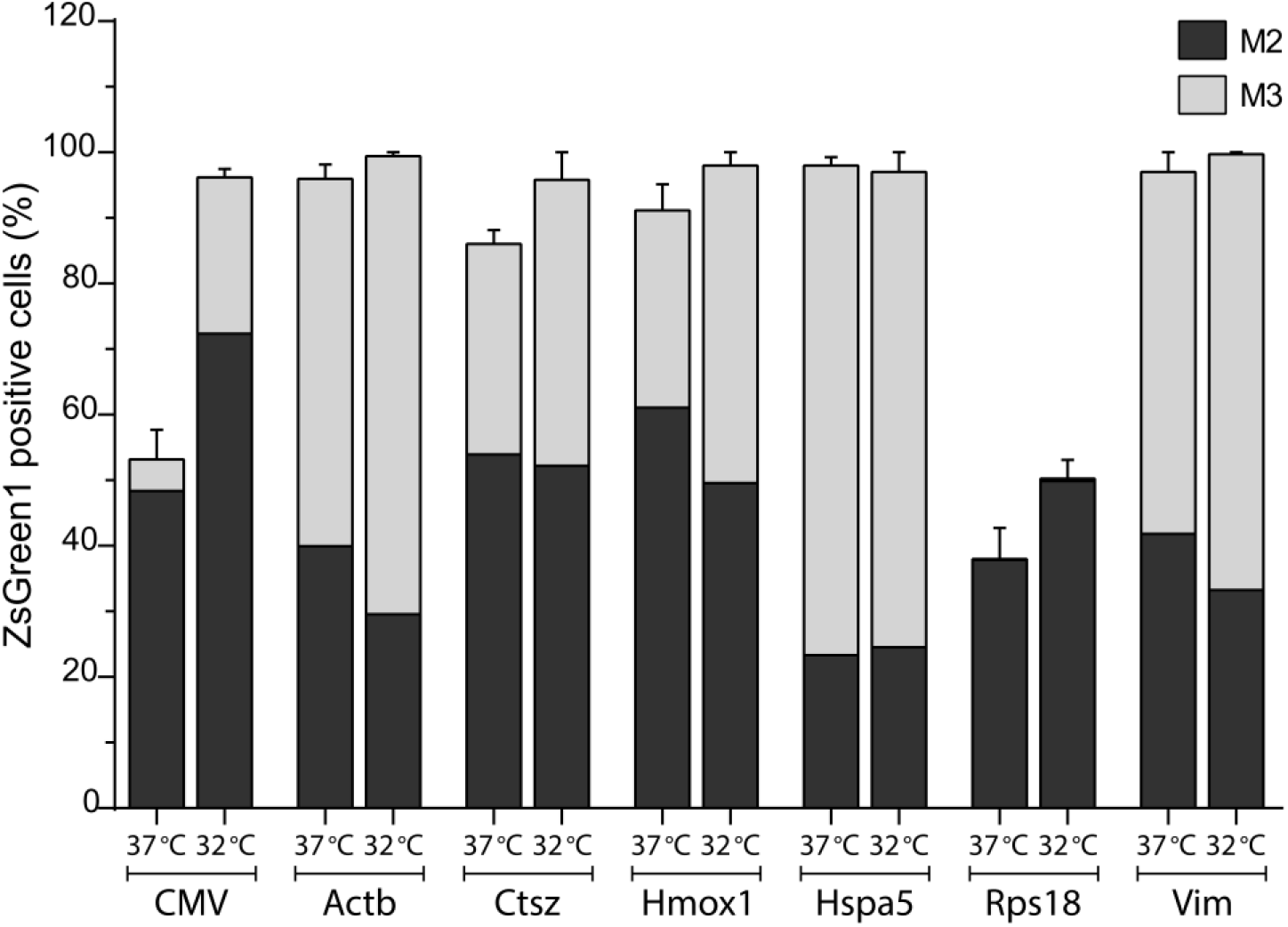
Impact of temperature shift on stable ZsGreen1 expression driven by endogenous promoters and CMV in CHO-K1 cells, during ten days upon cultivation at 37 °C or 32 °C. Bars show % of positive cells expressing moderate (black) and high (gray) levels of ZsGreen1 from samples taken on days 3, 5, 7 and 10. **M2:** positive cells with moderate fluorescence intensity, from 10^1^ to 10^3^ FAU (Fluorescence arbitrary units). **M3:** positive cells with high fluorescence intensity, FAU values higher than 10^3^.

In particular, CMV exhibited a marked increase in total ZsGreen1 positive cells at lower temperature (1.80-fold). However, it is important to consider that this promoter showed expression instability during several generations (Figure 2.A), which makes it difficult to ensure that such increment is caused by the temperature reduction. Interestingly, stable gene expression driven by different endogenous promoters remained relatively constant or even increased after a reduction of the culture temperature from 37 °C to 32 °C. Considering that candidate promoters (except for Rps18) showed expression stability during 45 generations (Figure 2.A), this is a promising indication of their potential to be used during biphasic cultures for the industrial production of recombinant proteins.

### 3.3. Characterization of endogenous promoters by *in silico* analysis of their sequences

Several factors contribute to the activity of a promoter, such as genomic cis-acting sequences, type of vector, cell line and sequence-specific transcription factors (TFs). These TFs are important regulators of biological processes that work by binding to transcriptional regulatory regions, such as promoters, to control the expression of their target genes. Each TF usually recognizes a collection of similar DNA sequences known as transcription factor binding sites (TFBSs), which can be represented as motifs using models like position weight matrices. ^[52]^ In addition, AT or GC rich sequences are often located in gene promoters and play a key role in transcription initiation. Characterization of TFBSs and other functional sequences is an important first step in understanding promoter activity, thus we applied *in silico* tools for a more comprehensive description of endogenous promoters, including prediction of the TSS, TATA box and putative TFBSs (Table 1). All promoters contain several putative specificity protein 1 (Sp1) binding sites, and some of them have other common motifs such as GC-box, E-box and AP-1. In 4 out of 6 endogenous promoters (Actb, Hspa5, Rps18 and Vim), a putative TATA-binding protein factor (TBP)-binding site (TATA box) was identified at an expected distance from the TSS.

**Table 1.** Potential functional promoter elements predicted by *in silico* analysis.

| Promoter | Size (bp) | TATA box | Distance TATA to TSS | %GC | CpG islands | Sp1 | GC-box | E-box | AP-1 | Total putative TFBS<sup>a</sup> |
| --- | --- | --- | --- | --- | --- | --- | --- | --- | --- | --- |
| **Actb** | 999 | Yes | 11 bp | 55 | - | 12x | 2x | 3x | - | 31 |
| **Ctsz** | 972 | No | - | 56 | - | 12x | 3x | 5x | - | 34 |
| **Hmox1** | 900 | No | - | 51 | - | 5x | 2x | 6x | - | 23 |
| **Hspa5** | 906 | Yes | 31 bp | 60 | 1x | 17x | 8x | 6x | 1x | 35 |
| **Rps18** | 1043 | Yes | 42 bp | 49 | - | 5x | - | 5x | 2x | 25 |
| **Vim** | 961 | Yes | 17 bp | 58 | - | 24x | 2x | - | 4x | 36 |
| **CMV<sup>b</sup>** | 1101 | Yes | 374 bp | 46 | 1x | 9x | 2x | 5x | 2x | 59 |

<sup>a)</sup> TFBSs were predicted using Alibaba2 online software.
<sup>b)</sup> This sequence contains the human cytomegalovirus (CMV) immediate-early enhancer/promoter region and a chimeric intron.

Most mammalian promoters (around 60% of mouse and human promoters) contain CpG islands, where methylation status of CG dinucleotides and their open chromatin organization can work as transcriptional regulators. ^[53]^ Hartl *et al*. (2019) ^[54]^ revealed that promoters having CpG islands show higher transcriptional activity than non CpG island promoters, and they tend to be active in different cell types. Also, they proposed that high CG content in promoter sequences might support transcriptional activity in an indirect manner, by increasing DNA accessibility so facilitating TFs binding. In this study, a putative CpG island region was found only in Hspa5 endogenous promoter, and in CMV promoter. Furthermore, GC content of endogenous promoters is relatively high, ranging from 49 to 60%, compared to 46% for CMV promoter and the average GC content of 42% of CHO-K1 genome. ^[55]^

We also found that endogenous promoters have numerous potential TFBSs (between 23 and 36 sites). Comparatively, CMV has the highest amount of putative TFBSs. However, no consistent pattern of TFBSs in endogenous promoter sequences that characterizes their transcriptional activities was detected. Sp1 was found several times in all promoters. Transcription factor AP-2 alpha A and regulator C/EBP alpha were identified one or two times in 5 out of 6 endogenous promoters. Nuclear factor 1 (NF1) binding motif was identified between 1 or 2 times in 4 out of 6 endogenous promoters. Particularly, CMV has 4 putative binding sites for C/EBP alpha and 7 for NF1. These TFs were analyzed for their expression on RNA-seq data. ^[9]^ NF1 was differentially expressed (Benjamini–Hochberg adjusted p-value <0.05), showing higher expression during stationary phase. AP-2 alpha A has a moderately constant expression during exponential and stationary stages, and Sp1 expression slightly decreased in stationary phase.

We also examined sequence conservation of endogenous CHO-K1 promoters by comparison to mouse, rat and human orthologous genes (Figure S5). As expected, higher values of sequence similarity were observed between promoters of CHO-K1, rat and mouse. In the particular case of Actb, Hspa5 and Vim promoters, sequence similarity ranges between 70 and 80%. Hmox1 and Rps18 promoters showed weak correlation between species while Ctsz exhibited an intermediate correlation, between 60 and 70%. These results could explain why in transient transfection assays endogenous CHO-K1 promoters exhibited higher activity in BHK-21 cells than in HEK293T cells. Moreover, Actb, Hspa5 and Vim promoters, which showed high correlation between species, are among the strongest promoters in CHO-K1, BHK-21 and HEK293T cells.

## 4. Discussion

For the industrial protein production a high rate of transgene expression is desired and hence strong viral-derived promoters are commonly used. ^[6], [56]^ Constitutive strong overexpression of the transgene under the control of such promoters, e.g. CMV, can cause a significant stress on the cell, upregulating the unfolded protein response and even inducing premature apoptosis. ^[20]^ Additionally, CMV is susceptible to epigenetic silencing due to DNA methylation and/or histone modifications, which is linked to production instability. ^[11]–[13], [18]^ Endogenous promoters, which function in a coordinated manner with cellular and bioprocess characteristics could help to avoid such drawbacks and, at the same time, maintain high expression levels of the transgene.

In this study, promoters of Actb, Ctsz, Hmox1, Hspa5, Vim and Rps18 genes were selected and isolated from CHO-K1 genome for the generation of new expression vectors. Their smaller size (around 1000 bp), compared to other CHO-K1 endogenous promoters previously described, ^[24]^ constitutes an important advantage. Transient transfections demonstrated that these endogenous promoters exhibited transcriptional activity not only in CHO-K1 cells but also in BHK-21 and HEK293T cell lines. Even though other works have identified CHO endogenous promoters as potential candidates for delivering recombinant gene expression, ^[21], [23]^ here we proved their potentiality not only in CHO but also in other commonly used cell lines.

Not only the level of transcription, but also its long-term stability is critical for recombinant protein production. Only a few studies of endogenous promoters have shown results of stable cell line generation; ^[19], [23]^ however, no analysis with respect to expression stability over extended cultivation time was performed. The analysis of CHO-K1 stable cell lines during several generations that was performed in this work revealed that ZsGreen1 expression driven by endogenous promoters was markedly higher compared to CMV, which contains the immediate-early enhancer/promoter region and a chimeric intron. In BHK-21 cells, endogenous promoters exhibited less expression stability and non-uniform transcriptional activity. Particularly, Hspa5 promoter demonstrated to be a proper candidate for CHO and BHK cell line generation. Temperature reduction is commonly used during production in the biopharmaceutical industry to improve protein production and extend cultivation time. Some reports have studied cold-shock responsive promoters upon temperature shift in transient transfections. ^[21], [22]^ Here we analyzed ZsGreen1 stable expression driven by endogenous promoters and CMV, after a reduction of CHO-K1 culture temperature from 37 °C to 32 °C. Temperature shift was beneficial for endogenous promoter activity since it improved significantly the protein productivity, which highlights the potential use of these promoters for industrial production. Mammalian promoters have a highly complex structure, not completely understood yet. Despite this great diversity, common features of all selected endogenous promoters were identified by *in silico* analysis, including high GC content, putative TFBSs such as Sp1, GC-box, E-box, AP-1. Hspa5, one of the strongest endogenous promoters, that exhibited high transcriptional activity for transient and also stable expression in all cell lines evaluated, contains a potential CpG island region and numerous TFBSs, which may explain its high transcriptional activity. Additionally, Hspa5 promoter revealed a strong sequence correlation between species (mouse, rat and human). It was observed also for other endogenous promoter candidates, such as Vim and Actb. Both promoters demonstrated high transcriptional activity in CHO-K1 and BHK-21 cells, but expression was stable only in the former one.

The endogenous promoters characterized in this work were strategically identified using the RNA-Seq data of CHO-K1 cells cultured under typical bioprocess conditions for recombinant protein production (high density followed by a temperature gradient), in order to search for the ones with high transcriptional activity under such conditions. These endogenous promoters constitute a promising alternative to CMV promoter due to their high strength, long-term expression stability and integration into the regulatory network of the host cell. These promoters may also comprise an initial panel for designing cell engineering strategies and synthetic promoters, as well as for industrial cell line development.

## Abbreviations

Actb: Actin beta
ANOVA: Analysis of variance
AP-1: Activator protein 1
AP-2 alpha: Activating enhancer binding protein 2 alpha
|BHK: Baby hamster kidney
bp: base pair
C/EBP alpha: CCAAT/enhancer binding protein alpha
CHO: Chinese hamster ovary
CpG: 5′-cytosine-phosphate-guanine-3′
Ctsz: Cathepsin Z
DMEM: Dulbecco’s Modified Eagle Medium
E-box: Enhancer box
EF1-alpha: Elongation factor 1-alpha
ELISA: Enzyme-Linked ImmunoSorbent Assay
FAU: fluorescence arbitrary units
FBS: fetal bovine serum
FPKM: Fragments per kilo base per million mapped reads
hCMV: Human cytomegalovirus
HEK: Human embryonic kidney
Hmox1: Heme oxygenase 1
Hspa5: Heat shock protein family A (Hsp70) member 5
IFNα: interferon alpha
MEM: Minimum Essential Medium
MFI: mean fluorescence intensity
NEO: Neomycin
NF1: nuclear factor 1
pBKS: pBluescript II KS plasmid
PCR: Polymerase chain reaction
RNA-seq: RNA sequencing
Rps18: Ribosomal protein S18
SP1: Specificity protein 1
SV40: Simian virus 40
TBP: TATA-binding protein
TF: Transcription factor
TFBS: Transcription factor binding site
TSS: Transcription start site
Vim: Vimentin
ZsGreen1: Zoanthus sp. green fluorescent protein.

## 4. Data availability

The data that support the findings of this study are available from the corresponding author upon reasonable request.

