## Supplemental Information for "Screening of CHO-K1 endogenous promoters for expressing recombinant proteins in mammalian cell cultures"

### Supporting Information

**Table S1.** Subset of genes that showed high transcriptional activity in both exponential and stationary phases of CHO-K1 suspension-adapted cells cultured in 1 L bioreactors.

| Gene | Day 7 Average FPKM | Day 7 SD | Day 8 Average FPKM | Day 8 SD | Day 14 Average FPKM | Day 14 SD | Day 15 Average FPKM | Day 15 SD |
|---|---|---|---|---|---|---|---|---|
| **COX3** | 3925.86 | 760.04 | 3027.07 | 91.39 | 4248.41 | 486.37 | 2850.89 | 359.77 |
| **S100a6** | 3544.64 | 396.05 | 3457.66 | 307.98 | 5865.75 | 76.51 | 6034.24 | 103.87 |
| **Actb** | 3139.92 | 80.63 | 3203.99 | 65.83 | 2900.21 | 9.65 | 3114.64 | 84.65 |
| **COX1** | 2779.44 | 1123.40 | 2017.05 | 338.67 | 6241.47 | 108.38 | 2853.56 | 1549.79 |
| **COX2** | 2402.68 | 673.22 | 1792.82 | 17.59 | 3598.01 | 370.89 | 2043.02 | 461.92 |
| **ND4** | 2277.26 | 692.19 | 1857.17 | 90.44 | 3390.58 | 331.77 | 1919.13 | 613.36 |
| **Fth1** | 2021.84 | 54.93 | 2303.08 | 78.13 | 2997.26 | 524.11 | 3324.25 | 996.32 |
| **S100a5** | 2013.52 | 229.29 | 1893.91 | 202.59 | 3239.22 | 50.02 | 3247.83 | 91.27 |
| **ATP6** | 1995.19 | 674.69 | 1427.15 | 170.71 | 2788.24 | 325.21 | 1398.10 | 443.51 |
| **Lgals1** | 1764.21 | 106.17 | 1694.33 | 157.44 | 2141.09 | 116.55 | 2213.97 | 207.12 |
| **CYTB** | 1565.36 | 532.93 | 1137.24 | 107.43 | 2058.71 | 198.86 | 1135.07 | 385.55 |
| **ND1** | 1445.35 | 609.80 | 883.17 | 123.52 | 2236.14 | 208.71 | 1031.94 | 440.50 |
| **ATP8** | 1324.66 | 513.19 | 830.31 | 79.43 | 1700.78 | 92.26 | 841.79 | 350.34 |
| **ND2** | 1285.64 | 547.80 | 861.05 | 131.34 | 1483.46 | 201.05 | 756.24 | 247.22 |
| **Gnas** | 1264.20 | 30.52 | 1232.28 | 72.40 | 880.74 | 98.10 | 948.86 | 129.98 |
| **Calr** | 1232.92 | 63.35 | 1216.82 | 82.47 | 1138.21 | 110.27 | 1199.07 | 102.07 |
| **ND4L** | 1183.90 | 426.65 | 925.93 | 27.81 | 1613.87 | 94.82 | 994.65 | 411.03 |
| **Cd63** | 1156.08 | 63.96 | 1091.16 | 41.15 | 1067.84 | 165.78 | 1139.28 | 233.35 |
| **Lgals3** | 1151.70 | 104.13 | 1178.41 | 51.11 | 1083.79 | 139.21 | 1106.60 | 288.71 |
| **Hspa5** | 1004.48 | 57.91 | 999.92 | 69.91 | 921.72 | 59.74 | 955.36 | 46.29 |
| **Ybx1** | 949.86 | 8.00 | 921.50 | 21.06 | 974.10 | 72.19 | 1009.47 | 41.75 |
| **Ctsz** | 903.98 | 85.58 | 908.52 | 28.13 | 875.97 | 145.38 | 945.17 | 181.53 |
| **Slc44a2** | 900.29 | 483.26 | 1121.91 | 282.45 | 878.75 | 78.19 | 592.37 | 63.96 |
| **Rps11** | 895.83 | 56.91 | 848.41 | 31.56 | 852.39 | 99.13 | 883.03 | 35.62 |
| **Myo9a** | 870.92 | 154.83 | 810.33 | 69.33 | 736.05 | 11.84 | 799.06 | 4.37 |
| **Kcmf1** | 852.56 | 45.01 | 659.27 | 28.68 | 636.71 | 121.21 | 622.51 | 44.75 |
| **Rps7** | 843.08 | 11.76 | 762.28 | 9.78 | 798.99 | 124.18 | 840.63 | 35.24 |
| **Psap** | 839.10 | 35.70 | 835.01 | 11.14 | 742.00 | 114.97 | 777.75 | 147.82 |
| **Rps18** | 817.61 | 79.35 | 782.18 | 16.12 | 822.96 | 158.70 | 877.34 | 137.48 |
| **Gapdh** | 816.34 | 73.87 | 734.94 | 33.89 | 563.16 | 1.35 | 634.81 | 37.82 |
| **Anxa2** | 787.96 | 54.64 | 825.50 | 33.77 | 1081.34 | 77.74 | 1151.70 | 42.61 |
| **Eef1a1** | 774.52 | 31.84 | 728.16 | 33.08 | 631.48 | 76.69 | 648.46 | 53.00 |
| **Bsg** | 774.49 | 58.18 | 722.36 | 5.58 | 690.24 | 91.32 | 754.67 | 166.54 |
| **Bub1** | 771.11 | 322.95 | 970.68 | 236.58 | 598.28 | 91.19 | 427.19 | 19.29 |
| **Mgst1** | 768.33 | 53.05 | 840.31 | 67.75 | 914.90 | 52.96 | 953.57 | 130.72 |
| **P4hb** | 768.16 | 55.71 | 779.49 | 27.16 | 820.62 | 72.77 | 873.55 | 78.41 |
| **Vim** | 745.32 | 71.04 | 832.77 | 106.50 | 1541.97 | 207.87 | 1599.93 | 202.25 |
| **Gtf2i** | 742.88 | 298.62 | 861.06 | 210.50 | 615.44 | 107.85 | 424.53 | 10.28 |
| **Ctsb** | 740.20 | 58.19 | 772.07 | 10.56 | 832.39 | 152.10 | 904.50 | 204.14 |

| | | | | | | | | |
|---|---|---|---|---|---|---|---|---|
| **Grn** | 733.70 | 37.16 | 788.65 | 23.30 | 1003.00 | 153.80 | 1126.35 | 255.27 |
| **Eif3j** | 718.87 | 36.04 | 694.35 | 43.26 | 707.96 | 178.52 | 721.63 | 229.97 |
| **S100a11** | 717.35 | 33.77 | 691.14 | 38.58 | 708.23 | 2.33 | 783.97 | 91.18 |
| **Atp5j2** | 715.67 | 26.77 | 627.52 | 34.75 | 622.15 | 13.10 | 677.80 | 29.82 |
| **Tom1** | 713.48 | 78.71 | 786.63 | 13.57 | 947.43 | 157.79 | 986.50 | 296.82 |
| **Hmox1** | 701.49 | 78.36 | 786.87 | 7.76 | 931.09 | 143.71 | 994.69 | 285.44 |
| **Tinagl1** | 695.38 | 78.43 | 739.73 | 33.15 | 1021.36 | 117.50 | 1120.93 | 145.50 |
| **Dazap1** | 693.37 | 60.35 | 637.56 | 32.00 | 607.42 | 97.22 | 610.78 | 84.28 |
| **Tpm4** | 691.12 | 46.47 | 674.98 | 31.80 | 440.21 | 5.57 | 432.93 | 45.27 |
| **Rpl38** | 681.19 | 40.98 | 631.90 | 19.62 | 669.84 | 5.57 | 667.74 | 24.67 |

**Table S2.** List of primers designed for PCR amplification of CHO-K1 candidate promoters. Restriction sites introduced at the 5’ ends of primers are not shown.

| Gene | Forward primer | Reverse primer |
|---|---|---|
| **Actb** | 5'-GGGAGATCTTCTCTGGTGGA-3' | 5'-GGGTTTTATAGGACGCCACA-3' |
| **Ctsz** | 5'-CCCCAGATTTGCCTCCGAG-3' | 5'-CGACCCTTTGACCTGACCC-3' |
| **Hmox1** | 5'-TAGAAGGGAGGCCTGGCAT-3' | 5'-CCGAGCCAGCCCTTTAAGT-3' |
| **Hspa5** | 5'-CGGGAACATTATGGGGCGA-3' | 5'-GGTCCTCCGGTCACTGTTG-3' |
| **Rps18** | 5'-GGGCAGACATAGGGAGTGC-3' | 5'-CATATCTCCGGCCCACACC-3' |
| **Vim** | 5'-ATGTTTGCGGTCTGGGAGG-3' | 5'-CAAGAGTGGCAGAGGACCG-3' |

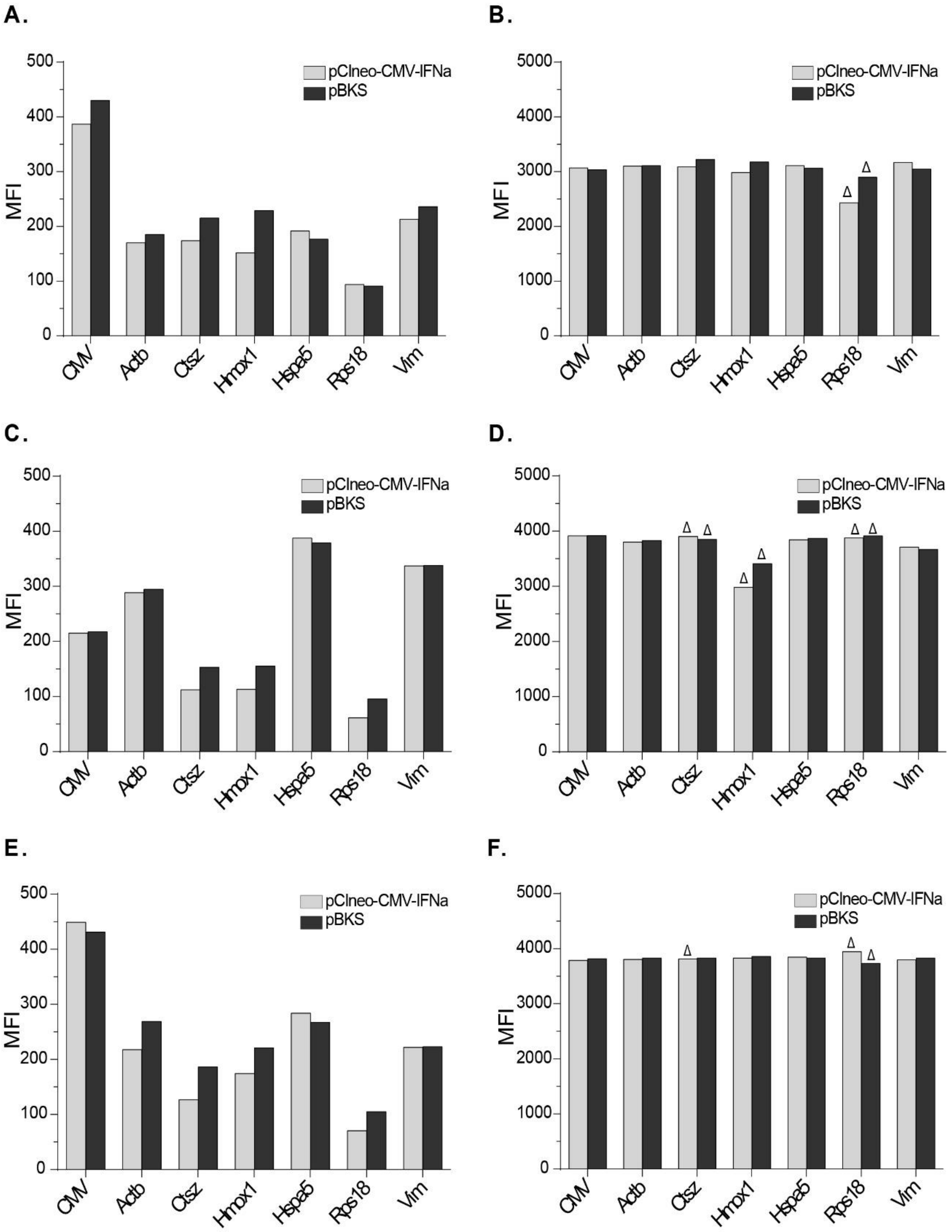


**Figure S1.** Effect of different promoters on transient transgene expression. pZsGreen1-1 vectors containing CMV and endogenous promoters were co-transfected with pBKS or pCIneo-CMV-IFNα plasmids into CHO, HEK293T and BHK-21 cells. **(A, C, E)** Moderate (M2) and **(B, D, F)** High (M3) MFI values of ZsGreen1 for CHO-K1, HEK293T and BHK-21 cells, respectively, in both co-transfection assays. **M2:** positive cells with moderate fluorescence intensity, from $10^1$ to $10^3$ FAU (Fluorescence arbitrary units). **M3:** positive cells with high fluorescence intensity, FAU values higher than $10^3$. Delta (**Δ**) indicates cell population in marker M3 less than 2%.

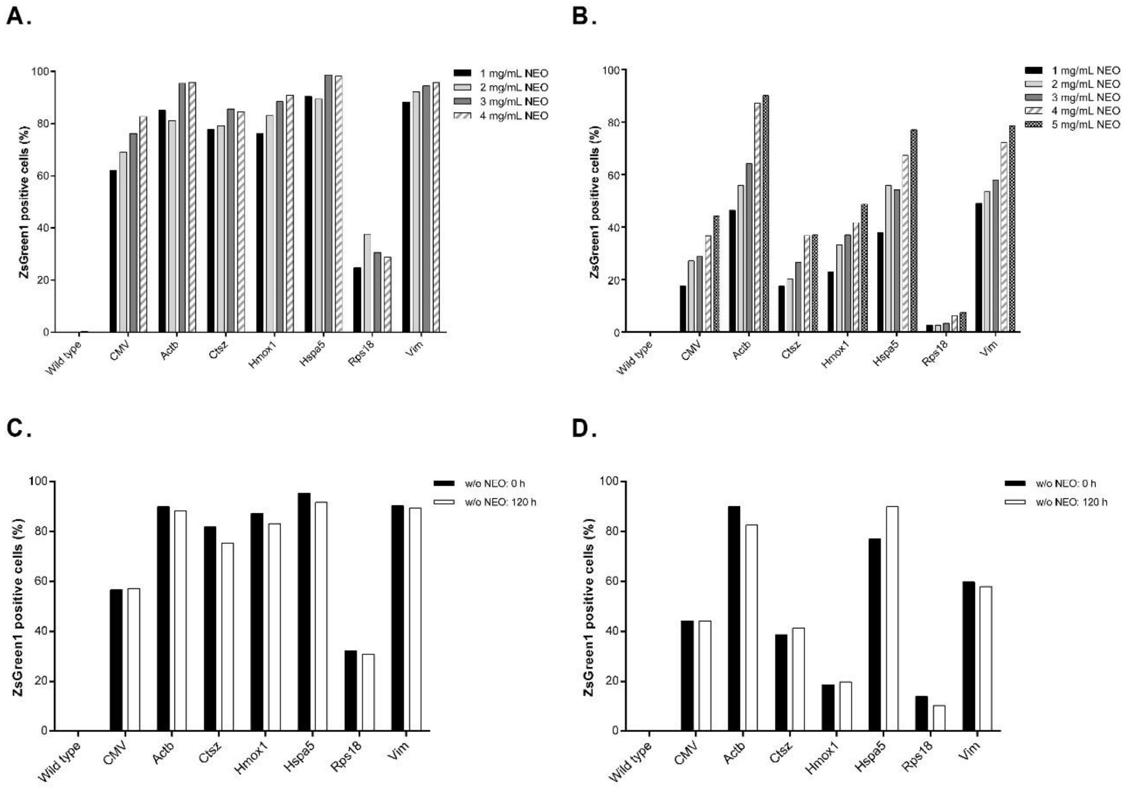


**Figure S2.** Generation of stable cell lines. **(A)** CHO-K1 cells cultured under neomycin (NEO) selection pressure from 1 to 4 mg/mL. **(B)** BHK-21 cells cultured under NEO selection pressure from 1 to 5 mg/mL. **(C)** ZsGreen1 expression levels of CHO-K1 cells growing with (4 mg/mL) and without NEO after 120 h. **(D)** ZsGreen1 expression levels of BHK-21 cells growing with (5 mg/mL) and without NEO after 120 h.

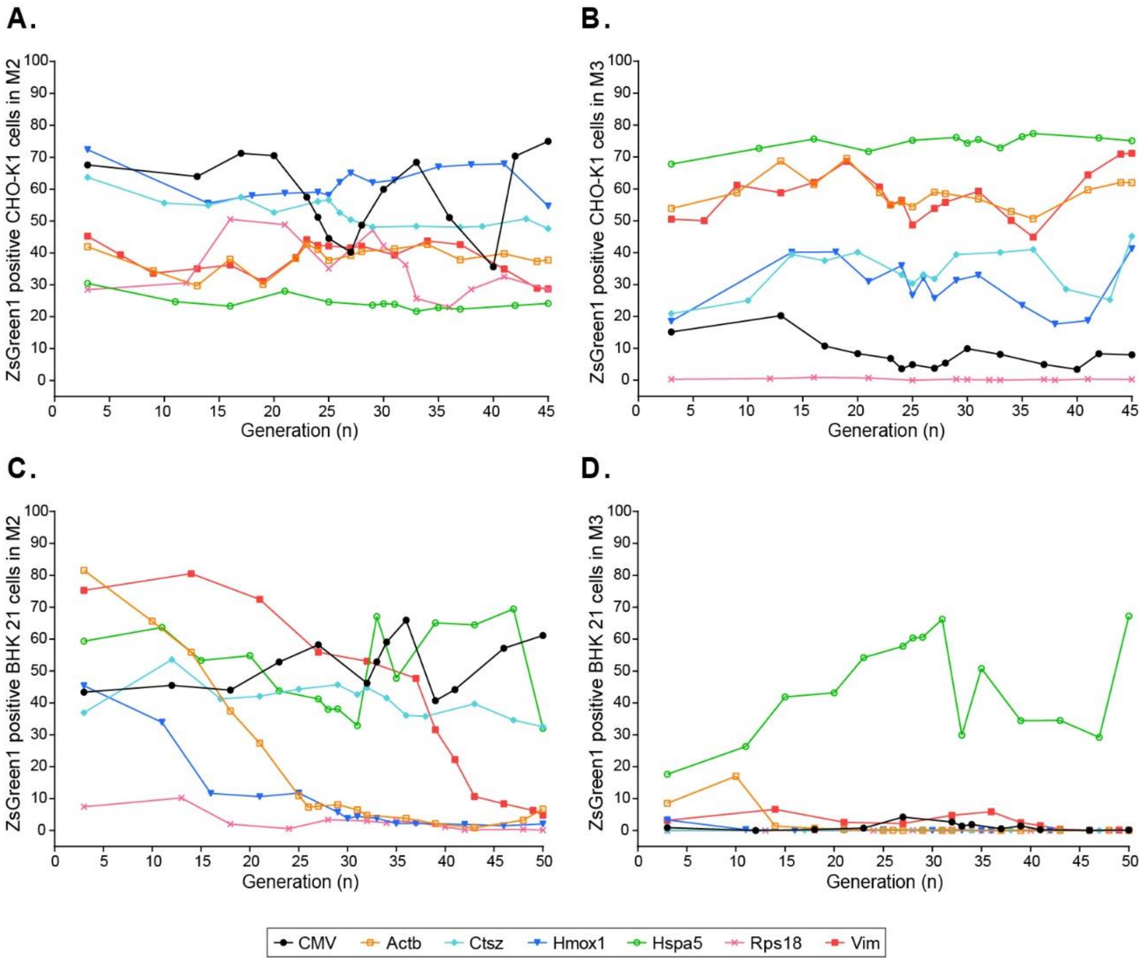


**Figure S3.** ZsGreen1 expression stability in CHO-K1 and BHK-21 cell lines grown until 45 and 50 generations, respectively, in the absence of neomycin selection pressure. ZsGreen1 positive CHO-K1 cells in marker M2 **(A)** and marker M3 **(B).** ZsGreen1 positive BHK-21 cells in marker M2 **(C)** and marker M3 **(D)**. **M2:** positive cells with moderate fluorescence intensity, from $10^1$ to $10^3$ FAU (Fluorescence arbitrary units). **M3:** positive cells with high fluorescence intensity, FAU values higher than $10^3$.

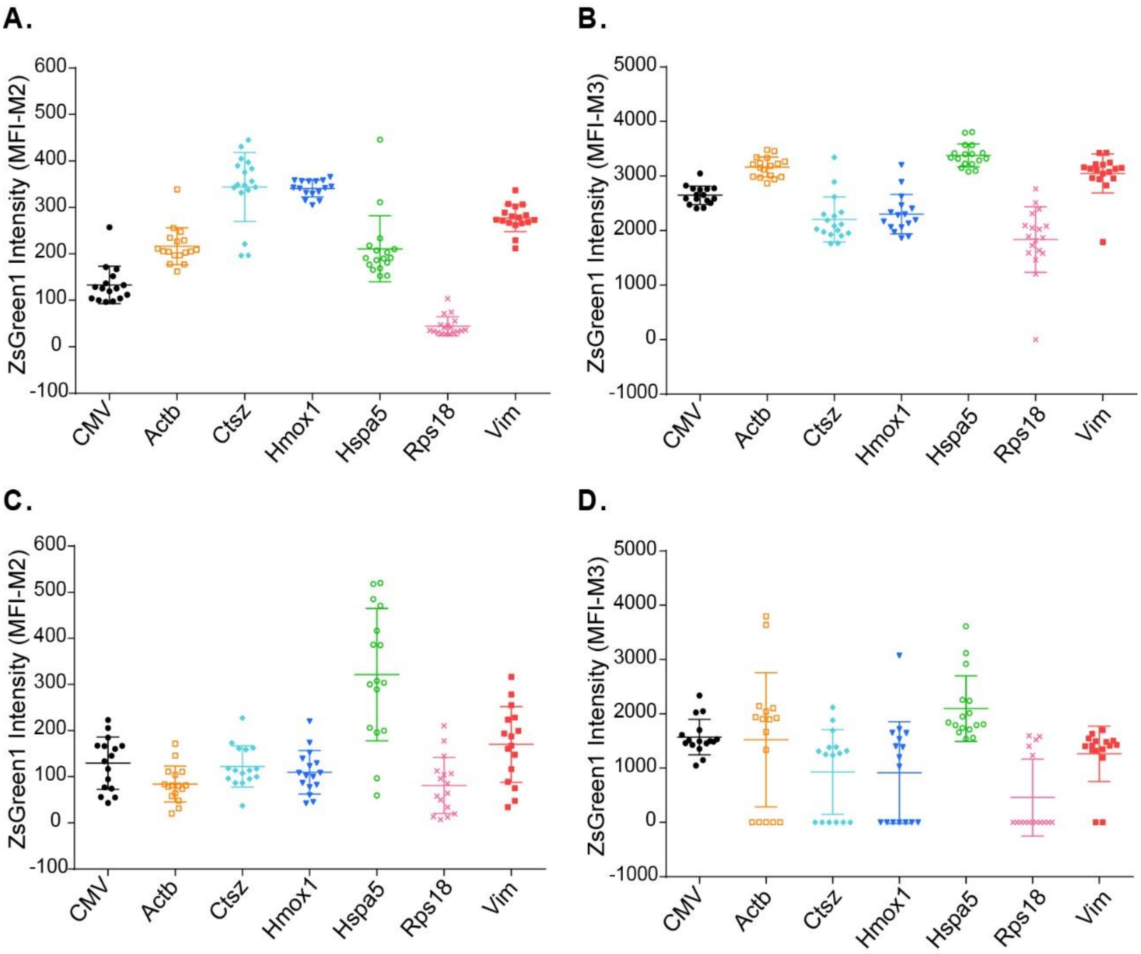


**Figure S4.** Mean Fluorescence Intensity (MFI) of ZsGreen1 expression under the control of endogenous promoters and CMV in CHO-K1 and BHK-21 cells during 45 and 50 generations, respectively. Moderate M2 **(A)** and high M3 **(B)** MFI values of ZsGreen1 for CHO-K1 cells. Moderate M2 **(C)** and high M3 **(D)** MFI values of ZsGreen1 for BHK-21 cells. **M2:** positive cells with moderate fluorescence intensity, from $10^1$ to $10^3$ FAU (Fluorescence arbitrary units). **M3:** positive cells with high fluorescence intensity, FAU values higher than $10^3$.

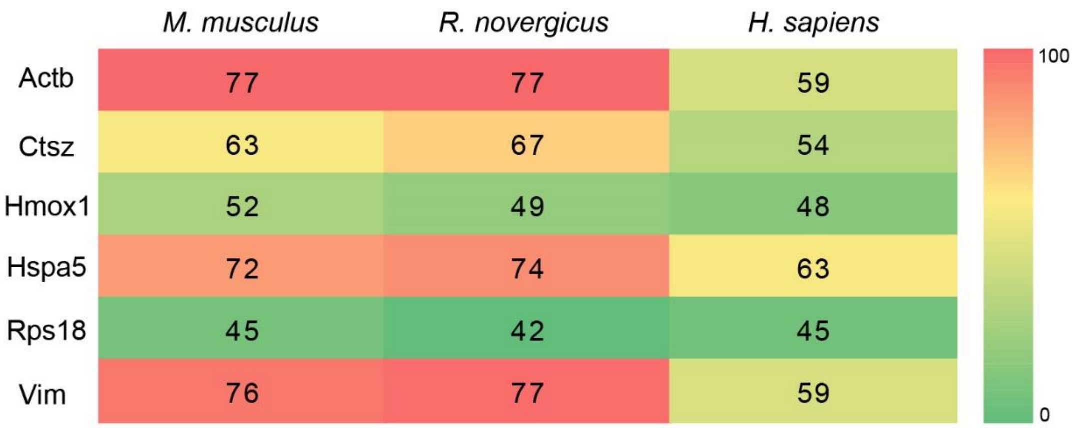


**Figure S5.** Sequence conservation of endogenous CHO-K1 promoters compared to mouse (*M. musculus*), rat (*R. novergicus*) and human (*H. sapiens*) orthologous sequences by pairwise global sequence alignment using the EMBOSS needle software. Percentages of similarity between promoter sequences are shown. Color bar scale represents low (green) and high (red) conservation.
